# Parallel maximal common subgraphs with labels for molecular biology

**DOI:** 10.1101/2024.05.10.593525

**Authors:** Wilfried Agbeto, Camille Coti, Vladimir Reinharz

## Abstract

Advances in graph algorithmics have allowed in-depth study of many natural objects from molecular biology or chemistry to social networks. Particularly in molecular biology and cheminformatics, understanding complex structures by identifying conserved sub-structures is a key milestone towards the artificial design of novel components with specific functions. Given a dataset of structures, we are interested in identifying all maximum common connected partial subgraphs between each pair of graphs, a task notoriously NP-Hard.

In this work, we present parallel algorithms over shared and distributed memory to enumerate all maximal connected common sub-graphs between pairs of arbitrary multi-directed graphs with labels on their edges. We offer an implementation of these methods and evaluate their performance on the non-redundant dataset of all known RNA 3D structures. We show that we can compute the exact results in a reasonable time for each pairwise comparison while taking into account a much more diverse set of interactions—resulting in much denser graphs—resulting in an order of magnitude more conserved modules. All code is available at https://gitlab.info.uqam.ca/cbe/pasigraph and results in the branch `results`.

## 1 Introduction

Graphs are increasingly being applied in various fields such as mathematics, biology, chemistry, and social network analysis. For instance, in biology, the tertiary structure of RNA can be represented as a graph, where each vertex represents a nucleotide identified by its base name and sequence number, and edges represent interactions between nucleotides, labeled by their interaction type. In this RNA graph model, searching for RNA structural motifs (which are recurring substructures appearing at non-homologous locations in one or more RNA molecules) is equivalent to searching for maximal common connected partial subgraphs (MCCPS).

The problem of finding the MCCPS between two graphs is a generalization that includes the connectivity aspect of the Maximum Common Partial Sub-graph (MCPS) problem, which is limited to finding only the largest maximal common partial subgraph between two graphs and is recognized in the literature as being NP-hard : for each vertex in the first graph, we find every similar vertex in the second graph. Edges of the graph are labeled with the type of interaction between the two nucleotides modeled by the graph. For each of these similar pairs, we try to extend the similar subgraph by comparing the edges connecting their neighbors, and we continue this extension as long as we find similar neighbors. Because of the computation-intensive nature of this problem, parallel computing is an attractive approach to speed-up the computation by taking advantage of the computation and memory capabilities of current high-performance computing platforms. Indeed, finding all the MCCPS requires an exhaustive exploration of the search space, and therefore has a high combinatorial complexity.

However, finding all the MCCPS between two graphs in parallel faces multiple challenges. The MCCPS are tied with each other: extending a subgraph can have consequences on other subgraphs that might be computed on other processes. Moreover, MCCPS computation times are very unbalanced. Therefore, computing them in parallel raises non-trivial load-balancing and inter-process communication challenges. Moreover, the number of solutions of an MCCPS problem can be exponential in the number of nodes, henceforth making exhaustive search highly computation-intensive.

In this paper, we present parallel algorithms to compute the MCCPS over shared memory, and distributed memory (hybrid approach). Our implementation works on multi-directed graphs with labeled edges; we present a performance evaluation on the dataset of all known RNA 3D structures. A limitation of previous works was the number of labels taken into account, the most efficient algorithms would fail on a computer with 8TB of RAM when all contacts between two large molecules were taken into account. The present algorithms easily overcome this limitation requiring only 186GB of RAM to process the entire dataset.

## 2 Related works

The problem of finding a Maximum Common Subgraph (MCS) has been studied for half a century. The problem of finding an MCS is divided into two categories of problems: MCIS (Maximum Common Induced Subgraph) [17,21,27] is the problem of finding a common induced subgraph with the largest number of nodes, and MCPS (Maximum Common Partial Subgraph) [2,22] is the problem of finding a common partial subgraph with the largest number of edges. Another distinction can be made between the connected case and the disconnected case. [9,8] have demonstrated that this problem is one of the most challenging in terms of algorithmic complexity.

There are two main types of exact approaches to solving this problem: the reduction approach to the maximum clique problem and the enumeration approach of common subgraphs (branch and bound) [15]. The reduction approach to the maximum clique problem relies on reformulating the problem into a compatibility graph. In this graph, each clique corresponds to a common subgraph. Thus, finding a maximum common subgraph is equivalent to identifying a maximum clique in the compatibility graph. In the enumeration approach of common sub-graphs, a search tree is constructed to explore all possible common subgraphs. The creation of subgraphs is halted when the algorithm determines that a subgraph cannot produce a solution larger than the best one found so far. The efficiency of these algorithms depends on the use of pruning heuristics to quickly remove unnecessary branches from the search tree [15]. The second approach is more efficient for small or sparse graphs. In all other cases, the first approach is more efficient [4].

Approximate [23] and parallel [13,6] approaches have also been proposed. Parallel approaches decompose the problem into independent sub-problems that can be solved simultaneously using domain partitioning. The MCS problem is one of the most difficult problems to parallelize due to its complex combinatorial nature and the interdependence among sub-problems. The subgraphs considered by the computation can span over multiple subdomains, leading to non-trivial communication patterns and load-balancing challenges.

Such irregular parallel applications can be dealt with using task-based programming with automatic load balancing, such as what was done for parameter space exploration. The parameter space is partitioned from points called *reference points*. From a reference point, polyhedraa round it are computed. Different polyhedra can be computed in parallel as their computation is independent of each other. However, it is impossible to know in advance which points will be included in the same polyhedron, so it is important to avoid computing the same polyhedron on different computational resources. This problem can therefore be connected to the MCS problem, in which the computation time to find a common subgraph and how the domain can be decomposed cannot be known before the common subgraph has been found. Heuristics and dynamic algorithms have been designed and evaluated in [1]. Their conclusion is that for small parameter spaces, a random distribution of points works well, and for larger parameter spaces, a dynamic domain decomposition with work stealing works better. [5] address a comparable parameter space partitioning problem. They compare different approaches, and the one that performs the best involves detecting and stopping redundant computations early and computing another task instead.

In this work, we will apply MCPS to the task of finding RNA motifs. An RNA molecule can be described as a sequence of nucleotides *ω* ϵ {*ACGU*}^*n*^, of a length from 10s to thousands of letters. They adopt 3D conformations through contacts (strengthened by hydrogen bonds) between the different nucleotides. The structure is known to form hierarchically [29], first canonical and wobble base pairs (A–U, G–C, and G–U contacts in the cis Watson–Crick/Watson– Crick `cWW` conformations) agregate into rigid *stems*, ladders of cWW contacts. These have experimentally determined energies [16] and are well captured by an efficient thermodynamic model in *𝒪* (*n*^3^), with the length of the sequence |*ω*| = *n* below 300 [11]. Yet the global configuration and binding geometries depend on finer conformations in the loops, positions outside of the *stems*, for which no energetic parameter are known. These new geometries can conveniently be described as graphs, where the nucleotides are nodes, the edges indicating contacts between the nodes. The geometry of these contacts is classified into 12 families by Leontis–Westhof [10]. They can be of type `cWW` as others depicted in Table 2. The backbone connects the nucleotides sequentially.

**Table 1.**
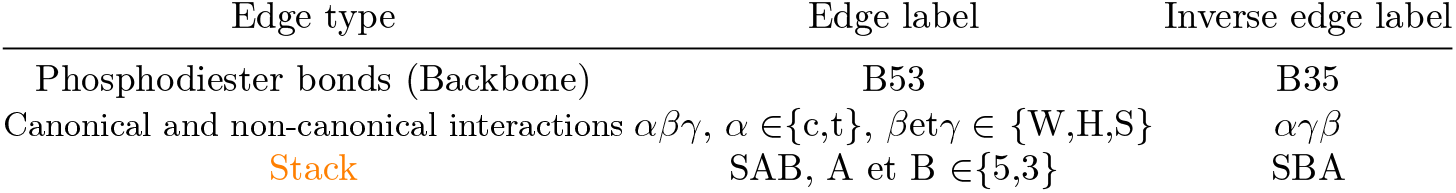
Labeling an RNA graph. cWW is canonical, other non-canonical.

Uncovering the main motifs, conserved sub-graphs, is a main challenge in the field necessary to support better tools for structure prediction as the design of artificial sequences [14]. Approaches directly from 3D structure have been limited to simple loops due to the complexity of atoms-to-atoms comparisons [19]. Alternatively, graph-based algorithms have been developed and have allowed to find motifs spanning multiple loops and with hundreds of nodes and edges [7,3,24,28]. Yet stackings, a core feature of RNAs, are too numerous to be analyzed with current methods. To overcome this task, we develop a memory-efficient distributed algorithm that allows to exhaustively enumerate all MCPPS when considering backbones, canonical and noncanonical contacts, as stackings.

## 3 Algorithms

In this section, we are presenting parallel algorithms to solve the MCCPS problem. We are presenting the sequential algorithm and data structures our parallel algorithms are based on (Section 3.1), then parallel algorithms over shared memory (Section 3.2) and distributed memory (Section 3.3).

### 3.1 Algorithm for Maximal Common Connected Partial Subgraphs

Let *g* and *h* be two directed graphs with edge labels. The approach used to find all MCCPS between *g* and *h* involves starting from an edge matching between *g* and *h* and adding neighboring edge matchings to it, considering all possibilities until reaching the maximal one. This approach is made of two steps: 1) Search for all edge matchings and 2) Extension of edge matchings.

*Search for all edge matchings (Algorithm 1)* Two edges in *g* and in *h* correspond together if they have the same label and direction. We use a list describing the edge types involved to treat an edge matching and its reverse edge matching as a single edge matching. For example, for edgeTypeList=[{CWS,CSW}, {TWW,TWW}], the first element of edgeTypeList implies that if there exists an edge (*a, b, CWS*) in *g*, then there also exists an inverse edge (*b, a, CSW*) in *g*. The function `SearchAllEdgeMatchings` returns the list of all found edge matchings (`edgeMatchList`), as well as two other lists: **edgeMatchIdList1** contains the identifier of the edge in *g* for each edge matching, and **edge-MatchIdList2** contains the identifier of the edge in *h* for each edge matching.

*Extension of edge matchings* Edge matchings are the elementary patterns constituting the MCCPS (A MCCPS can be represented as a set of edge matchings between *g* and *h*.). In this section, we present the algorithm that extends an edge matching to find all MCCPS containing it.

Extending an edge matching **s** involves adding neighboring edge matchings to it through a breadth-first search (BFS) traversal until maximization. Maximization is achieved when there are no more neighbors to explore.

During the extension of an edge matching, conflict events may occur. A conflict event happens when a neighboring edge matching cannot be added to the CCPS (Common Connected Partial Subgraph: subgraph under construction and not yet maximal). This occurs when a neighboring edge matching has a node ag|ah that cannot be added to the CCPS, as the latter either has a node *a*^*′*^*g*|*ah* or a node *ag*|*a*^*′*^*h*, or both (with *a*^*′*^*g* ≠ *ag* and *a*^*′*^*h* ≠ *ah*).

For each conflict, we create a new extension branch containing a new CCPS to be extended. To obtain the new CCPS, we create a copy of the MCCPS by removing the nodes and edges in conflict with the conflicting edge matching and adding the conflicting edge matching to the copy. Since the extension of an edge matching **s** should yield all MCCPS containing **s**, any conflicts that imply removing **s** are ignored.

Extending an edge matching **s**, combined with conflict management, allows us to find all MCCPS containing **s. (1)**Conflict management enables the extension algorithm to backtrack and explore alternatives to find additional MCCPS. However, multiple conflicts can lead to the same MCCPS, posing a challenge. To mitigate this, before creating a new extension branch, we check if the new CCPS to be extended is already included in any previously found MCCPS starting from **s**. If so, the branch is not created. This inclusion check involves searching for an exact subgraph isomorphism that preserves edge labels and node identifiers.

**(2)**Since extending an edge matching **s** yields all MCCPS containing **s**, then the extension of multiple edge matchings can yield identical MCCPS. To address this issue, when an edge matching **e** can be added to a CCPS resulting from extending an edge matching *s*_*i*_, then we check if **e ϵ {***s*_0_, *s*_1_, *s*_2_, …, *s*_*i*−1_}, where *s*_*i*_ is the edge matching at index *i* in **edgeMatchList**. If this is the case, the edge matching is not added to the CCPS. The subgraph obtained at the end of extending the CCPS may not be an MCCPS, meaning it may not be maximal. If at least one of the edge matchings **e**, which were ignored by the CCPS, can be added to the obtained subgraph, then it is not maximal and therefore not considered as an MCCPS.

Before describing the parallel algorithms, it is important to present the different types of tasks.

**Task of extension an edge matching:** This task involves extending an edge matching. Thanks to (2), these tasks can be executed in parallel independently without communication or synchronization. **Branch extension task:** These tasks arise from conflict resolution. Due to (1), they can be executed in parallel, but they need to synchronize to avoid searching for an already found MCCPS. It is also important to note that conflict resolution is local to the extension of a given edge matching.

Our objective is to simultaneously (if possible) reduce idle time (improving workload balancing) and the synchronization costs of execution units.

### 3.2. Shared Memory Algorithm

Algorithm 1 gives the parallel algorithm in the shared memory model. We base our approach on the **PCAM** method, which includes the following steps:

– **Partitioning:** The problem is decomposed into fine-grained tasks.
– **Communication:** Identification of dependencies between tasks.
– **Agglomeration:** Once the tasks and their dependencies have been determined in the previous steps, tasks can, if necessary, be combined into larger tasks to improve performance (data movements).
– **Mapping:** Distribution of tasks across processing units.

*Search for all edge matchings* The partitioning step is achieved using a one-dimension domain decomposition on the `edgeTypeList` to create one task for each element of `edgeTypeList`. The computation time for each of these tasks is irregular and can take advantage of a dynamic scheduling policy such as the one provided by OpenMP.

*Extension of edge matchings* The set of edge matchings also uses a one-dimensional domain decomposition to create one task for each edge matching. We obtain fine-grained tasks because extending edge matchings is strongly unbalanced between the tasks. In the following, we focus on a work-stealing approach to balance the workload automatically. We are using threads as processing units. We explored the impact of the scheduling strategy being used here, and the results are presented in Section 4.1.

*Dynamic Distribution* Each thread receives a task corresponding to the extension of an edge matching. When it is done, it moves on to another task. In this approach, tasks are independent, so it is not necessary to communicate or synchronize between tasks. However, this approach suffers from significant load imbalance when the distribution of the number of MCCPS per edge matching is highly unbalanced (in some practical cases, extending an edge matching took more than 80% of the total execution time).

#### Algorithm 1

SearchAllEdgeMatchings Parallel

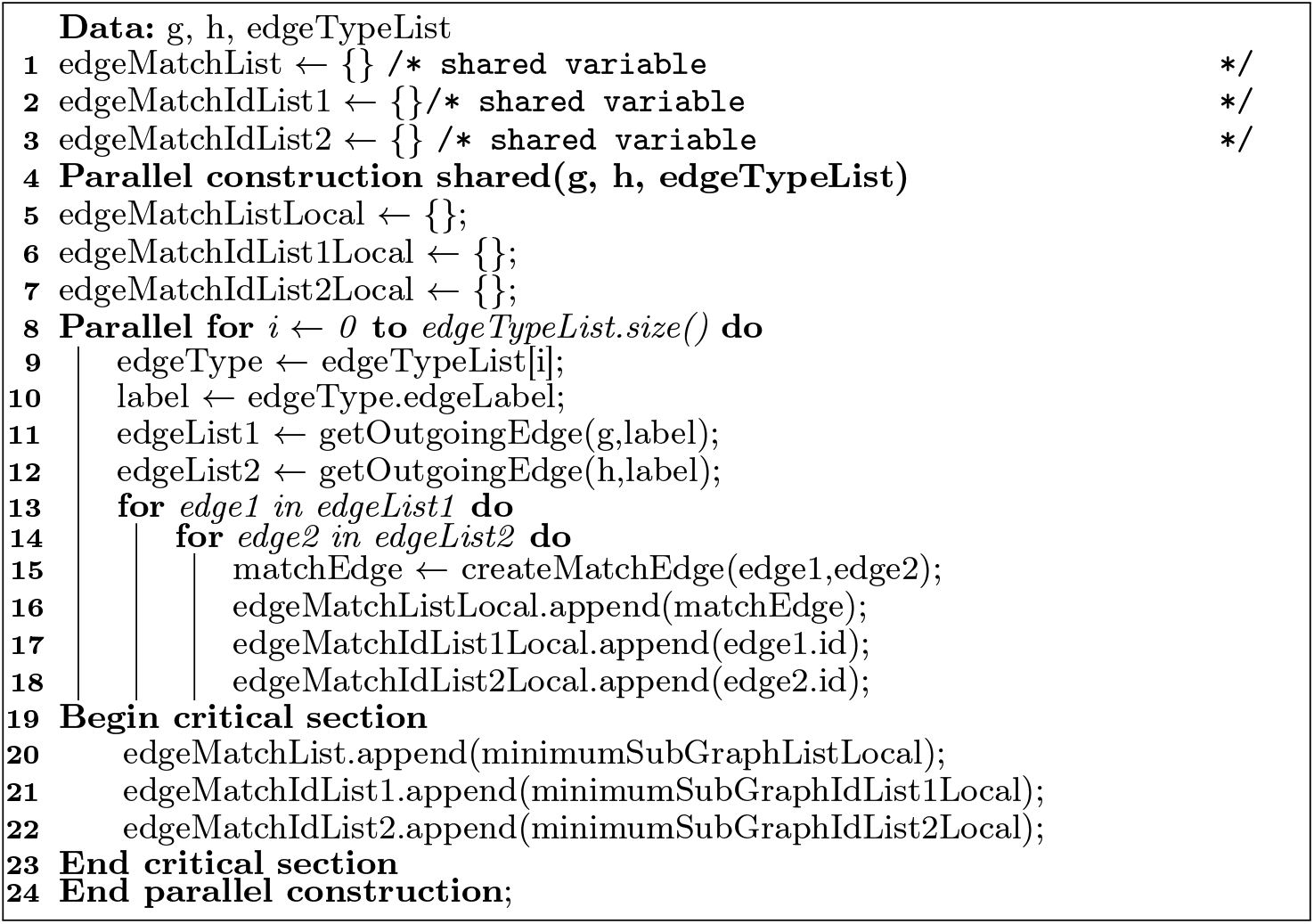

#### Nested

Threads are organized in groups. The master thread of each group receives a task corresponding to the extension of an edge matching, and the branch extension tasks resulting from conflict resolution are distributed among the threads of the same group. Threads within the same group must synchronize to avoid extending a branch that will yield an already-found MCCPS. When it is done, the group moves on to another task of extension an edge matching. To avoid duplications, a list containing the MCCPS already found is shared with exclusive access among the threads of the same group. The process to avoid duplication is described in Section 3.1. Although this method improves load distribution, complex tasks can monopolize a group of threads and unbalance the others, making it crucial to precisely adjust the size and number of groups to avoid underutilization of resources.

#### Work Stealing

This approach is similar to the dynamic distribution one, except that when there is no more tasks that extend an edge matching are exhausted, idle threads steal branch extension tasks from active threads. This approach allows for better workload balancing. To reduce task synchronization overhead, threads do not directly share their branch extension tasks; they only do so when at least one other thread is available. A thread knows if there is at least one other available thread through the shared variable *numThreadFree*. When there are no more tasks to extend edge matchings, idle threads set the value at index *rankThread* in *threadStateList* to 1, indicating they are available, and increment *numThreadFree* by 1. When at least one thread is available, only one of the busy threads shares its branch extension tasks: the thread with the smallest thread id using the function *isYourRound* that takes *threadStateList* and *rankThread* as input, traverses *threadStateList*, and returns 1 if it finds only 1, indicating that the calling thread should share its tasks. Otherwise, the function returns 0.

### 3.3 Distributed Memory Algorithm

*Search for All Edge Matchings* The starting point of the algorithm is edge matching. This can be distributed easily like on shared memory: every process starts from a set of labels and communication is necessary at this step. Each process executes the same algorithm for shared memory (Algorithm 1).

*Extension of Edge Matchings* Work stealing is more complex in a distributed environment because synchronization between branch extension tasks to avoid duplicates (searching for an already found MCCPS) would require a significant amount of communication between processes, making this approach inefficient. Therefore, we have proposed a hybrid approach where the tasks of extending an edge matching are shared among processes using the master-worker scheme, while the threads within the same group use the work stealing approach to manage the branch extension tasks. This helps avoid costly communications between processes in order to prevent duplicates.

With this method, there is a risk that some processes may be underutilized compared to others. However, by increasing the number of computing nodes, we enhance computational power compared to a shared-memory approach. Moreover, having more computing nodes can offset the underutilization of processes and thus improve overall efficiency.

### 3.4 From MCCPS to biological RNA motifs

A few constraints are necessary to transition from the mathematical object toa relevant biological motif. It is known that interesting RNA motifs have a cyclic structure, and each node is involved in a non-canonical interaction (interaction labeled neither `CWW` nor `B53`) [18,12].

As a postprocessing step, for each of the MCPPS, we iteratively remove nodes that do not fulfill the following conditions: (1) nodes must be connected by an interaction other than B53, (2) if two nodes *n*_1_, *n*_2_ are connected by a `cWW`, there must be a node *n*_3_ neighbor to *n*_1_ or *n*_2_ involved in a non-canonical or stacking interaction with a fourth node (i.e. we do not extend stacks of only canonical base pairs), and (3) each node belongs to a cycle.

It is clear that all maximal recurrent RNA motifs must belong to an MCPPS and be extractable from it, else another bigger RNA motif must contain it, a contradiction.

## 4 Performance evaluation

We implemented these algorithms using OpenMP (4.5) provided with gcc 11.4 and OpenMPI (4.1.2). The code was optimized by g++ using -O3. The performance data presented in this section was measured on Calcul Québec’s ma-chine Narval^3^. The set of 1722 non-redundant RNA structures was taken from RNA3DHub version 3.269 [20]. The graphs for each structure were built with annotations of the contact geometries from FR3D [26]. The smallest graph has two nodes and two edges, the biggest 16 770 nodes and 37 582 edges. Over 200 graphs have a node count greater than or equal to 100.

To validate the parallel constructions of MCCPS, we compared them to the sequential version of our algorithm and also compared the results with those from previous work [25,28].

### 4.1 Shared memory performance

First, we compared the performance of the three shared memory algorithms presented in Section 3. We picked two graphs randomly; one graph has 390 vertices and 1003 edges, and the other one has 1800 vertices and 4282 edges. The performance comparison is presented in Figure 2. We can see that all three algorithms benefit from the parallelization when a few threads are used, and beyond 4 threads, the performance of the nested and dynamic algorithms stalls whereas the work-stealing algorithm keeps scaling.

**Fig. 1.**
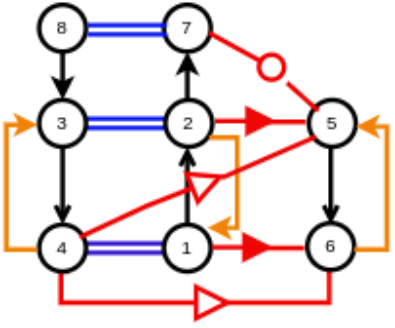
RNA graph, blue double lines are `cWW` interactions forming stems, black edges connect positions sequentially, red edges are between loops, orange edges are stacks.

**Fig. 2.**
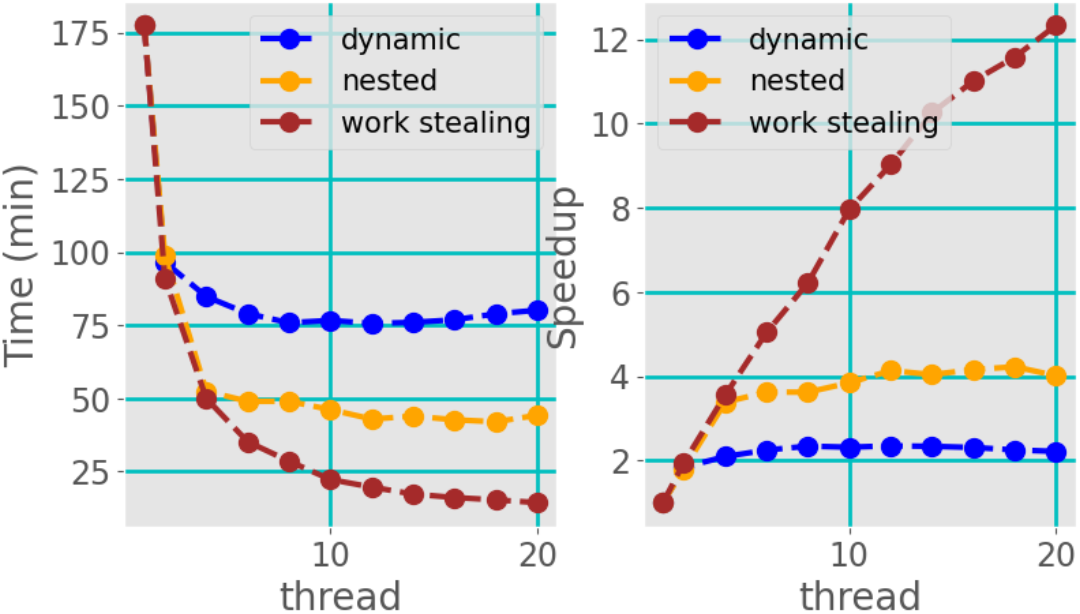
Scalability of the shared memory algorithms.

We picked randomly 7 pairs of graphs from the RNA3DHub dataset and we compared the performance obtained with the three algorithms on these graphs using 10 threads. As presented in Figure 3, the work-stealing approach was faster on every graph we used.

**Fig. 3.**
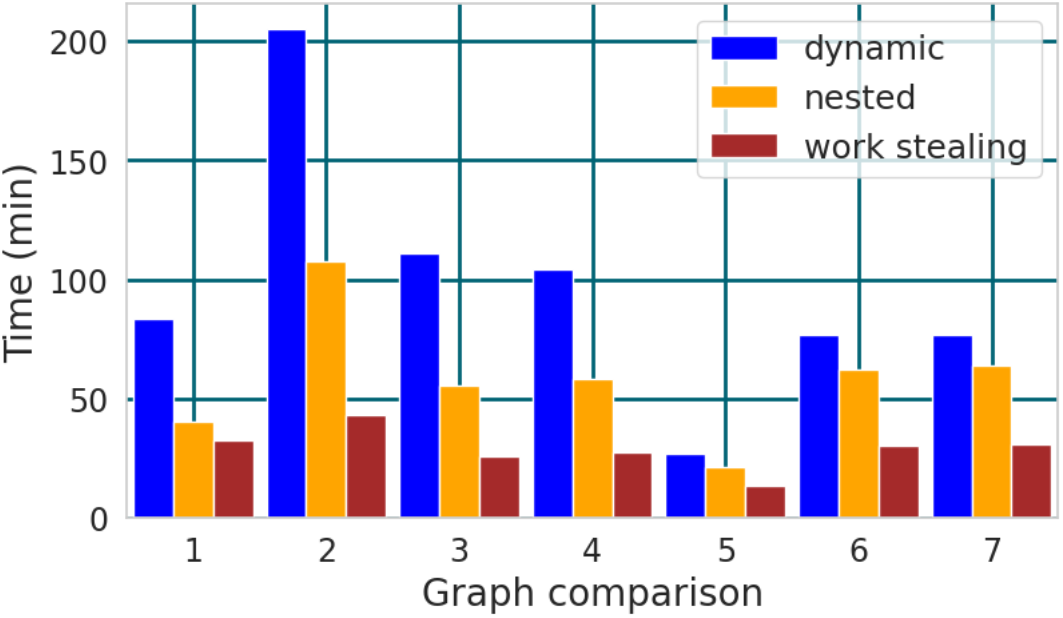
Performance comparison between the shared memory algorithms.

### 4.2 Hybrid algorithm

Current parallel architectures are hierarchical: they are made of multi-core nodes connected by a high-speed network. As a consequence, parallel programs can follow a hybrid approach and use shared-memory algorithms on multiple cores of the same node, and distributed-memory algorithms between nodes. Since distributed-memory algorithms can be used on several cores of a given node, the distribution of the processing units (processes and threads) between shared memory and distributed memory is a tuning parameter of the program.

In our case, we used a multi-thread algorithm over shared memory and multiple processes over distributed memory. We evaluated the impact of the threads *times* processes combination on the performance. Since Narval’s nodes feature 48 cores, we compared the performance obtained using one process and 48 threads per process, and all integer combinations up to 48 single-threaded processes.

Based on the observations made in Section 4.1, we used the work-stealing algorithm on the multi-thread part of the computation. The performance comparison on a single node is presented in Figure 4. The speed-up is indicated over every bar. We can see that the approach using 48 threads gives the best performance.

**Fig. 4.**
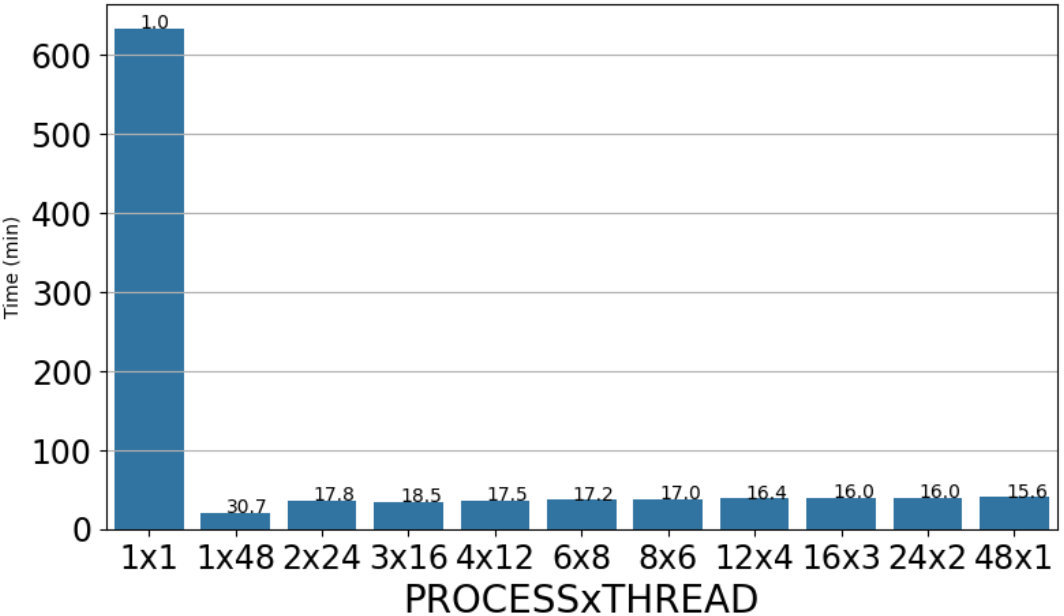
Hybrid algorithm: number of threads per process for a given number of cores. The two graphs compared have 1247 and 1742 nodes, and 3492 and 4412 edges, respectively

### 4.3 Performance of the hybrid algorithm

Figure 5 presents the scalability of the hybrid approach. We used all the cores of each node, with different combinations of the number of processes per node and the number of threads per process. As explained in the previous sections, we used the work-stealing approach. We can see that using one process per node and 48 threads per process gives the best performance, confirming the expectation presented in Section 4.2. The performance improves with the number of nodes up to 6 nodes (288 cores). This size was sufficient for us to obtain the results we needed: the aggregated memory of these 6 nodes was enough to perform the computation, and the execution time was reasonable.

**Fig. 5.**
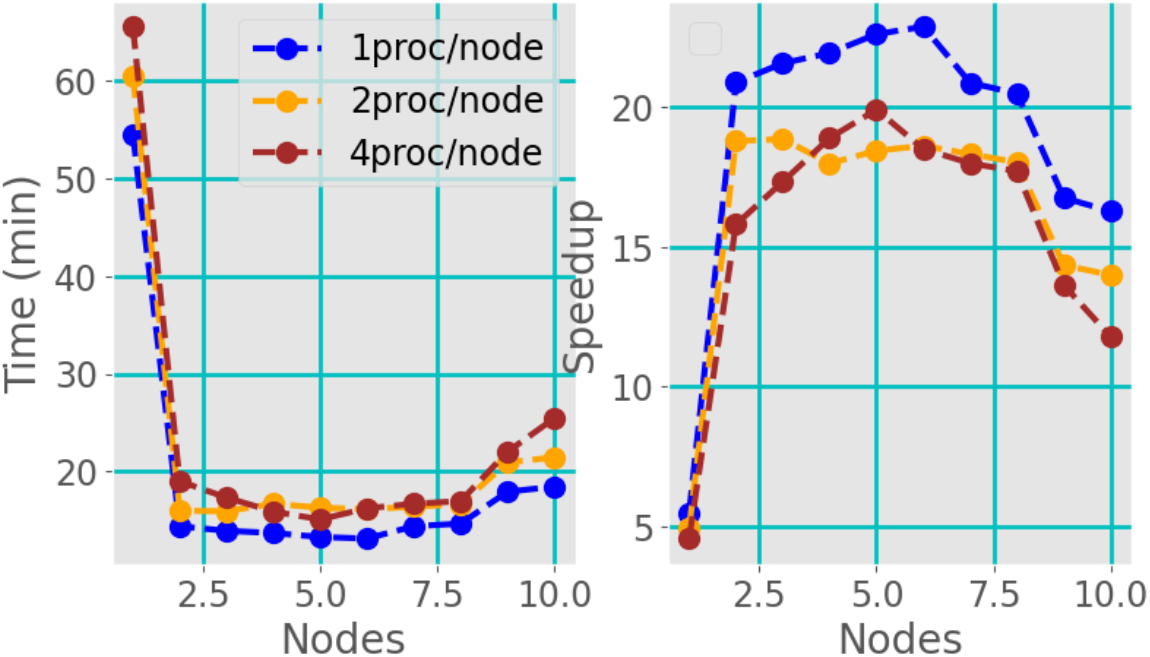
Scalability of the hybrid approach. The two graphs compared have 210 and 942 nodes, and 2910 and 7235 edges, respectively

We tried to speed up the computation more by using a larger number of cores, and the performance worsened. Thanks to the performance we reached with the distributed version, it was possible to complete the computation and get results for the bioinformatics community. However, getting results faster is always desirable, for instance to complete more computations in a given makespan. Performance analysis suggests when the number of processes is large enough, the workload is distributed between them over a few steps of the master-worker scheme, and the automatic load balancing mechanism cannot compensate for load unbalances between nodes. Future works will investigate other distributed approaches, for example with more dynamic load-balancing approaches.

### 4.4 RNAs

We used our parallel program to perform a pairwise comparison of all the non-redundant existing RNA structures to identify all the structural motifs they contain (see Sec. 3.4. It took 4 weeks on Narval, which is significantly faster than previous works [25,28]. It required a total of 186 GB of memory. It is important to note that the results obtained were not achievable with the methods proposed in previous work, even with 8TB of RAM. Additionally, we found a larger number of structural motifs. We found 157 344 structural motifs for a total of 209 750 474 occurrences. The smallest motif has 3 nodes and 6 edges, and the largest 1135 nodes and 5566 edges.

The A-minor motif is well known for its structural importance as a general tool to join loops, yet this versatility makes it hard to predict in practice. We show in Fig. 6 the first map of stacking variants of the A-minor motif. The numbers on the figure indicate motif coverage.

**Fig. 6.**
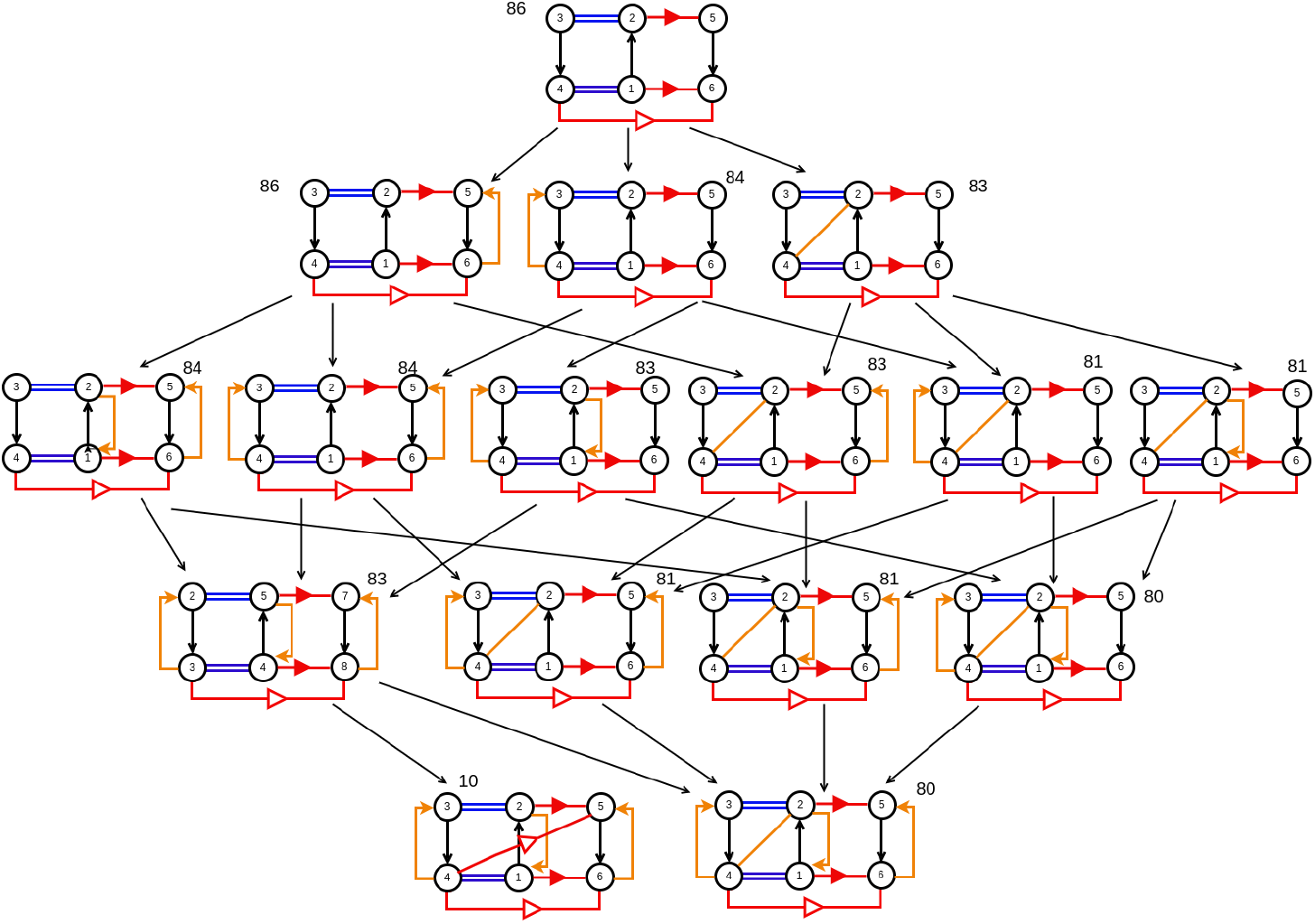
Variation of A-Minor motifs with stacking. The orange lines, oriented and unoriented, represent stacking interactions S53 and S55, respectively.

## 5 Conclusion

In this paper, we present novel parallel algorithms for the exhaustive enumeration of Maximum Common Partial Subgraphs between two arbitrary multi-directed graphs, admitting labels over edges. This problem has a high computation complexity, but its irregular nature makes it non-trivial to parallelize. We offer an open-source implementation in C and evaluate it using complex RNA structure graphs as benchmarks. We show that our algorithms scale over shared and distributed memory using multiple threads and multiple processes, and with efficient memory usage.

In molecular biology, MCPS can be indicative of important functional motifs. In the case of RNA structures that have an efficient graph representation, motifs finding algorithms always ignored different key features such as stackings, known to be important but too numerous to be fully taken into account, until now.

With the algorithms presented here, we can now exhaustively list all recurrent motifs creating novel maps of natural variants with stackings. Such information will be leveraged in structure prediction, molecular design, and binding target prediction.

The abundance and size of motifs raise new questions that will require efficient parallel algorithms to solve. Studying RNA structures is not a static endeavor, every week new structures emerge and have to be analyzed and merged with our dataset. With over 150 000 motifs now, their organization and clustering are another challenge. Annotating all their positions in the few thousands of known RNAs requires several terabytes of data. While the number of known RNA structures remains low, the amount of information to analyze has been increasing over the last decades and will continue to do so.

RNA structures are one limited example of what can be achieved now using the computation power offered by these parallel algorithms. Labels on nodes are currently ignored, but with new experimental techniques allowing annotation of chemical modifications in RNAs using nanopore sequencing, we can expect such a feature to be highly desirable.

## Footnotes

3 https://docs.alliancecan.ca/wiki/Narval/en

