## Supplementary material for "Parallel maximal common subgraphs with labels for molecular biology"

### 6 Supplementary material

Here we present our Algorithm for Maximal Common Connected Partial Subgraphs and its shared memory version in more detail.

#### 6.1 Algorithm for Maximal Common Connected Subgraphs (MCCPS)

Let $g$ and $h$ be two directed graphs with edge labels. The approach used to find all **MCCPS** between $g$ and $h$ consists of starting from an edge matching between $g$ and $h$ and adding neighboring edge matchings to it, considering all possibilities until reaching the maximal. This method consists of the following tasks:

1. Search for all edge matchings;
2. Extension of edge matchings;

**Search for All Edge matchings** An edge in $g$ and an edge in $h$ correspond together if they have the same label and the same direction. we use a list describing edge types to treat an edge matching and its reverse edge matching as a single edge matching. For example, for edgeTypeList = $[\{CWS, CSW\}, \{TWW, TWW\}]$, the first element of edgeTypeList implies that if there exists an edge $(a, b, \text{CWS})$ in $g$, then there also exists a reverse edge $(b, a, \text{CSW})$ in $h$.

```
Data: g, h, edgeTypeList
edgeMatchList ← {};
edgeMatchIdList1 ← {};
edgeMatchIdList2 ← {};
for i ← 0 to edgeTypeList.size() do
    edgeType ← edgeTypeList[i];
    label ← edgeType.edgeLabel;
    edgeList1 ← getOutgoingEdge(g,label);
    edgeList2 ← getOutgoingEdge(h,label);
    for edge1 in edgeList1 do
        for edge2 in edgeList2 do
            matchEdge ← createMatchEdge(edge1,edge2);
            edgeMatchList.append(matchEdge);
            edgeMatchIdList1.append(edge1.id);
            edgeMatchIdList2.append(edge2.id);
        end
    end
end
```

**Algorithm 2:** SearchAllEdgeMatches

The function SearchAllEdgeMatches (Algorithm 2) returns the list of all found edge matchings (**edgeMatchList**) as well as two other lists:

- **edgeMatchIdList1:** It contains the identifier of the edge in $g$ for each edge matchings.
- **edgeMatchIdList2:** It contains the identifier of the edge in $h$ for each edge matchings.

**Extension of Edge matchings** Edge matchings are the elementary patterns constituting the MCCPS. In this section, we present the algorithm for extending an edge matching to find all MCCPS containing it.

**Starting point: (Algorithm 3, Line 4)** A starting point **s** for two graphs $g$ and $h$ is an edge matching between $g$ and $h$. In other words, the set of starting points is the set of edge matching.

**Extension from a starting point s: (Algorithm 4)** This involves finding an MCCPS by performing a breadth-first search (BFS) from the starting point **s**. The subgraph is dynamically constructed by adding neighboring edge matchings to the starting point **s** until maximization is achieved. Maximization is reached when there are no more neighbors to explore. An edge matching is added to the ongoing subgraph only when it has a node that is already present in the subgraph, which is necessary to maintain connectivity.

**Common Connected Partial Subgraph (CCPS):** It's a connected subgraph under construction from a starting point **s** and is not yet maximal.

**Conflicts: (Algorithm 4 line 9; line 15-17)** During the extension from a starting point **s**, conflict events may occur. A conflict event happens when a neighboring edge matching cannot be added to the CCPS. This occurs when a neighboring edge matching has a node $a_g|a_h$ that cannot be added to the CCPS because the latter either has a node $a'_g|a_h$ or a node $a_g|a'_h$, or both (with $a'_g \neq a_g$ and $a'_h \neq a_h$). A conflict suggests the existence of a different MCCPS than the one being currently produced. Indeed, a conflict indicates the existence of an MCCPS containing at least one edge matching (the edge matching responsible for the conflict) that the current MCCPS under production will not contain. When a CCPS yields conflicts, we store for each conflict the edge matching (we store its index in **edgeMatchList**) responsible for it, and we defer the handling of these conflicts until the MCCPS is obtained.

**Conflict Management: (Algorithm 5)** When a CCPS yields an MCCPS, we handle all the conflicts it has generated by creating for each conflict a new extension branch containing a new CCPS to be extended. To obtain the new CCPS, we create a copy of the MCCPS by removing the nodes and edges conflicting with the conflicting edge matching and adding the conflicting edge matching to the copy. The obtained copy may contain multiple connected components; the other connected components are removed, and only the connected component containing the conflicting edge matching is retained. Two cases are possible:

- The connected component contains only the edge matching responsible for the conflict; in this case, the new extension branch is destroyed because it will

yield the same MCCPS as the extension with the edge matching responsible for the conflict as a starting point.
- The connected component does not contain only the edge matching responsible for the conflict; in this case, a new extension branch is created with the connected component.

During the extension from a starting point **s**, all found MCCPS must contain **s**, so conflicts involving nodes from **s** are ignored.

**(1)**Extension from a starting point **s**, combined with conflict management, allows finding all MCCPS containing **s**. Indeed, conflict management enables the extension algorithm to backtrack on its choices and make alternative ones to discover other MCCPS. However, a challenge with this approach is that multiple conflicts can lead to the same MCCPS. To address this, before creating a new extension branch, we check if the new CCPS to be extended is included in any of the MCCPS already found from the starting point **s**. If so, the branch is not created. For the inclusion check, we search for an exact subgraph isomorphism (preserving edge labels and node IDs) (Algorithm 6).

**(2)**Since extending from a starting point **s** yields all MCCPS containing **s**, extending from different starting points may result in identical MCCPS. To address this issue, when an edge matching **e** can be added to a CCPS resulting from extending from a starting point $s_i$, we check if $\mathbf{e} \in \{s_0, s_1, s_2, ..., s_{i-1}\}$, where $s_i$ is the edge matching at index i in **edgeMatchList**. If this is the case, the edge matching is not added to the CCPS (Algorithm 4 lines 10-12). The subgraph obtained at the end of extending the CCPS may not be an MCCPS, meaning it may not be maximal. If at least one of the edge matchings **e**, which were ignored by the CCPS, can be added to the obtained subgraph, then it is not maximal and therefore not considered as an MCCPS (Algorithm 3 lines 16-18).

**Normalized form of subgraphs:** The normalized form we use to represent subgraphs is an adjacency list. During the extension from a starting point **s**, new extension branches are created, each containing a new **CCPS** to extend. The issue is that the number of extension branches rapidly explodes, especially for large graph instances. Additionally, these branches are stored in memory waiting for extension, which consumes a lot of memory unnecessarily. To address this problem, we developed a compressed form to represent subgraphs.

**Compressed form of subgraphs:** As we saw previously, subgraphs are merely a set of edge matchings, so we can represent a subgraph by an array of integers (compressed form) containing the indices in **edgeMatchList** of the edge matches that compose it. The compressed form uses much less memory than the normalized form, with a ratio of approximately 10. During its lifecycle, a subgraph is represented in this compressed form except during the extension process. The conversion from compressed form to normalized form occurs in $\theta(n)$ time, where $n$ is the number of edges in the subgraph.

```
Data: edgeMatchList, edgeMatchIdList1, edgeMatchIdList2
MCCPSList ← {} /* List of all MCCPS */
for i ← 0 to edgeMatchList.size() do
    branches ← {};
    s ← edgeMatchList[i] /* starting point */
    branche ← createBranche();
    branche.graphCompressedForm.append(i);
    branches.append(branche);
    MCCPSListTmp ← {} /* list of MCCPS from starting point s */
    for branche in branches do
        cf ← branche.graphCompressedForm;
        if isIncluded(cf, MCCPSListTmp) ≠ 1 then
            branche.graphNormalForm ← getNormalForm(cf);
            result ← extendBranche(s, branche, edgeMatchList);
            conflictingMatchesEdge ← result[0];
            ignoredMatchesEdge ← result[1];
            if isMaximal(branche.graphCompressedForm, ignoredMatchesEdge)
              then
                MCCPSListTmp.append(branche.graphCompressedForm);
                conflictManagement(edgeMatchList, conflictingMatchesEdge,
                  branches, MCCPSListTmp, s, branche);
            end
        end
        free(branche);
    end
    MCCPSList.append(MCCPSListTmp);
end
```

**Algorithm 3:** ExtensionOfEdgeMatches

```
Data: s, branch, edgeMatchList
edgeMatchListTmp ← {};
conflictingMatchesEdge ← {};
ignoredMatchesEdge ← {};
for node in branch.graphNormalForm.nodes() do
    edgeMatchListTmp ← getEdgeMatchListContainNode(node) /* Returns
        edge matches that contain the node as a parameter */
    for edgeMatch in edgeMatchListTmp do
        id ← edgeMatch.index;
        if id not in branch.graphCompressedForm then
            if canBeAdd(branch.graphNormalForm, edgeMatch) then
                if s.index > id then
                    ignoredMatchesEdge.append(id);
                else
                    branch.graphCompressedForm.append(id);
                    branch.graphNormalForm.append(edgeMatch);
                end
            else
                if id not in conflictingMatchesEdge then
                    conflictingMatchesEdge.append(id);
                end
            end
        end
    end
end
```

**Algorithm 4:** extendBranch

```
Data: edgeMatchList, conflictingMatchesEdge, branches, MCCPSList, s,
      branche
for index in conflictingMatchesEdge do
    edgeMatch ← edgeMatchList[index];
    nf ← branch.graphNormalForm;
    nodes ← getConflictingNodes(nf);
    subgraph ← copyGraphWhithoutNodes(nf,nodes);
    subgraph.add(edgeMatch);
    subgraph ← removeComponent(subgraph, edgeMatch)/* Remove the
        other connected components and keep the connected component
        containing edgeMatch                                          */
    if subgraph.size() > 1 and s in subgraph then
        if isIncluded(subgraph, MCCPSList) ≠ 1 then
            newBranch ← createBranch();
            newBranch.graphCompressedForm ←
              getCompressedForm(subgraph);
            free(subgraph);
            branches.append(newBranch);
        end
    end
end
```

**Algorithm 5:** conflictManagement

```
Data: graphCompressedForm, MCCPSList
for MCCPS in MCCPSList do
    if graphCompressedForm.size() ≤ MCCPS.size() then
        if graphCompressedForm ⊆ MCCPS then
            return 1;
        end
    end
end
return 0;
```

**Algorithm 6:** isIncluded

The function isIncluded is used for exact subgraph isomorphism testing. By utilizing the compressed form (set of integers) of a subgraph, our algorithm has a complexity of $\Theta(n * m)$, where $n$ and $m$ are the number of edges in the two compared graphs. The isIncluded function is also used in the conflictManagement function because the number of MCCPS found between creating and extending a branch may differ.

##### 6.2 Shared Memory Algorithm

Before describing the parallel algorithms, it is important to present the different types of tasks.

- **Task of extension an edge matching:** This task involves extending an edge matching. Thanks to (2), these tasks can be executed in parallel independently without communication or synchronization.
- **Branch extension task:** These tasks arise from conflict resolution. Due to (1), they can be executed in parallel, but they need to synchronize to avoid searching for an already found MCCPS. It is also important to note that conflict resolution is local to the extension of a given edge matching.

Our objective is to simultaneously (if possible) reduce idle time (improving workload balancing) and the synchronization costs of execution units.

**Extension of edge matchings** We performed a one-dimensional domain decomposition on the set of starting points to have one task for each starting point. We used fine-grained tasks because the workload is highly unbalanced for extensions from a starting point. For distributing tasks across processing units, we employed various scheduling strategies to try to balance the workload.

*Dynamic Distribution* Each thread receives a task corresponding to the extension of an edge matching. When it is done, it moves on to another task. In this approach, tasks are independent, so it is not necessary to communicate or synchronize between tasks. However, this approach suffers from significant load imbalance when the distribution of the number of MCCPS per edge matching is highly unbalanced.

```
Data: edgeMatchList, edgeMatchIdList1, edgeMatchIdList2
MCCPSList ← {} /* shared variable */
Parallel construction shared(edgeMatchList, edgeMatchIdList1,
  edgeMatchIdList2);
MCCPSListLocal ← {};
parallel for i ← 0 to edgeMatchList.size() do
    branches ← {};
    s ← edgeMatchList[i]/* starting point */
    branche ← createBranche();
    branche.graphCompressedForm.append(i);
    branches.append(branche);
    MCCPSListTmp ← {} /* list of MCCPS from starting point s */
    for branche in branches do
        cf ← branche.graphCompressedForm;
        if isIncluded(cf, MCCPSListTmp) ≠ 1 then
            branche.graphNormalForm ← getNormalForm(cf);
            result ← extendBranche(s, branche, edgeMatchList);
            conflictingMatchesEdge ← result[0];
            ignoredMatchesEdge ← result[1];
            if isMaximal(branche.graphCompressedForm, ignoredMatchesEdge)
              then
                MCCPSListTmp.append(branche.graphCompressedForm);
                conflictManagement(edgeMatchList, conflictingMatchesEdge,
                  branches, MCCPSListTmp, s, branche);
            end
        end
        free(branche);
    MCCPSListLocal.append(MCCPSListTmp);
begin critical section;
    MCCPSList.append(MCCPSListLocal);
end critical section;
sameEdgePostProcessing(MCCPSList) /* Parallel */
End parallel construction;
```

**Algorithm 7:** ExtensionOfEdgeMatches parallel dynamic distribution

*Nested* Threads are organized in groups. The master thread of each group receives a task corresponding to the extension of an edge matching, and the branch extension tasks resulting from conflict resolution are distributed among the threads of the same group. Threads within the same group must synchronize to avoid extending a branch that will yield an already-found MCCPS. When it is done, the group moves on to another task.

```
Data: edgeMatchList, edgeMatchIdList1, edgeMa1.2chIdList2
MCCPSList ← {} /* shared variable */
Parallel construction shared(edgeMatchList, edgeMatchIdList1,
  edgeMatchIdList2);
MCCPSListLocal ←{};
parallel for i ← 0 to edgeMatchList.size() do
|   branches ← {};
|   s ← edgeMatchList[i]/* starting point */
|   branche ← createBranche();
|   branche.graphCompressedForm.append(i);
|   branches.append(branche);
|   MCCPSListTmp ← {} /* list of MCCPS from starting point s */
|   Parallel construction private(branche);
|   branchesSize ← 0;
|   branchesSizeTmp ← 0;
|   while branchesSizeTmp < branches.size() do
|   |   branchesSize ← branches.size();
|   |   parallel for j=branchesSizeTmp to branchesSize do
|   |   |   branche ← branches[j];
|   |   |   cf ← branche.graphCompressedForm;
|   |   |   if isIncluded(cf, MCCPSListTmp) ≠ 1 then
|   |   |   |   branche.graphNormalForm ← getNormalForm(cf);
|   |   |   |   result ← extendBranche(s, branche, edgeMatchList);
|   |   |   |   conflictingMatchesEdge ← result[0];
|   |   |   |   ignoredMatchesEdge ← result[1];
|   |   |   |   if isMaximal(branche.graphCompressedForm,
|   |   |   |     ignoredMatchesEdge) then
|   |   |   |   |   begin critical section;
|   |   |   |   |   MCCPSListTmp.append(branche.graphCompressedForm);
|   |   |   |   |   conflictManagement(edgeMatchList,
|   |   |   |   |     conflictingMatchesEdge, branches, MCCPSListTmp, s,
|   |   |   |   |     branche);
|   |   |   |   |   end critical section;
|   |   |   |   end
|   |   |   end
|   |   |   free(branche);
|   |   branchesSizeTmp ← branchesSize;
|   end
|   End parallel construction;
|   MCCPSListLocal.append(MCCPSListTmp);
begin critical section;
    MCCPSList.append(MCCPSListLocal);
end critical section;
sameEdgePostProcessing(MCCPSList) /* Parallel */
End parallel construction;
```

**Algorithm 8:** ExtensionOfEdgeMatches parallel nested version

*Work Stealing* This approach is similar to the dynamic distribution one, except that when there is no more tasks that extend an edge matching are exhausted, idle threads steal branch extension tasks from active threads. This approach allows for better workload balancing. To reduce task synchronization overhead, threads do not directly share their branch extension tasks; they only do so when at least one other thread is available(Algorithm 9, lines 19-22) . A thread knows if there is at least one other available thread through the shared variable *numThreadFree*. When there are no more tasks to extend edge matches, idle threads set the value at index *rankThread* in *threadStateList* to 1, indicating they are available, and increment *numThreadFree* by 1(Algorithm 9, lines 44-46) When at least one thread is available, only one of the busy threads shares its branch extension tasks: the thread with the smallest thread id using the function *isYourRound* that takes *threadStateList* and *rankThread* as input, traverses *threadStateList*, and returns 1 if it finds only 1, indicating that the calling thread should share its tasks. Otherwise, the function returns 0.

```
Data: edgeMatchList, edgeMatchIdList1, edgeMatchIdList2
MCCPSList ← {} /* shared variable */
threadStateList ← {} /* shared variable */
numThreadFree ← 0 /* shared variable */
Parallel construction shared(edgeMatchList, edgeMatchIdList1,
  edgeMatchIdList2);
MCCPSListLocal ← {};
shareTask ← 0;
Parallel for i ← 0 to edgeMatchList.size() do
    branches ← {};
    s ← edgeMatchList[i]/* starting point */
    branche ← createBranche();
    branche.graphCompressedForm.append(i);
    branches.append(branche);
    MCCPSListTmp ← {} /* list of MCCPS from starting point s */
    branchesSize ← 0;
    branchesSizeTmp ← 0;
    while branchesSizeTmp < branches.size() do
        branchesSize ← branches.size();
        for j=branchesSizeTmp to branchesSize do
            if not shareTask and numThreadFree>0 then
                if isYourRound(threadStateList,threadRank) then
                    shareTask ← 0;
                end
            end
            Do Task if(shareTask) private(branche);
            branche ← branches[j];
            cf ← branche.graphCompressedForm;
            if isIncluded(cf, MCCPSListTmp) ≠ 1 then
                branche.graphNormalForm ← getNormalForm(cf);
                result ← extendBranche(s, branche, edgeMatchList);
                conflictingMatchesEdge ← result[0];
                ignoredMatchesEdge ← result[1];
                if isMaximal(branche.graphCompressedForm,
                  ignoredMatchesEdge) then
                    begin critical section;
                    MCCPSListTmp.append(branche.graphCompressedForm);
                    conflictManagement(edgeMatchList,
                      conflictingMatchesEdge, branches, MCCPSListTmp, s,
                      branche);
                    end critical section;
                end
            end
            free(branche);
            End Task;
        if shareTask then
            Task Wait;
        end
        branchesSizeTmp ← branchesSize;
    end
    MCCPSListLocal.append(MCCPSListTmp);
Begin critical section;
    MCCPSList.append(MCCPSListLocal);
End critical section;
threadStateList[rankThread] ← 1;
Begin critical section;
    numThreadFree++;
End critical section;
sameEdgePostProcessing(MCCPSList) /* Parallel */
End parallel construction;
```

**Algorithm 9:** ExtensionOfEdgeMatches parallel work stealing version
